# STING controls T cell memory fitness during infection through T cell intrinsic and IDO dependent mechanisms

**DOI:** 10.1101/2022.03.19.484992

**Authors:** Michael J. Quaney, Rebecca J. Newth, Knudson M. Karin, Vikas Saxena, Curtis J. Pritzl, Chris S. Rae, Peter Lauer, Mark A. Daniels, Emma Teixeiro

## Abstract

Stimulator of interferon genes (STING) signaling has been extensively studied in inflammatory diseases and cancer while its role in T cell responses to infection is unclear. Using *Listeria monocytogenes* strains engineered to induce different levels of c-di-AMP, we found that strong STING signals impaired T cell memory upon infection via increased Bim levels and apoptosis. Unexpectedly, reduction of TCR signal strength or T cell-STING expression decreased Bim expression, T cell apoptosis and recovered T cell memory. We found that TCR signal intensity coupled STING signal strength to the Unfolded Protein Response (UPR) and T cell survival. Under strong STING signaling, IDO inhibition also reduced apoptosis and led to a recovery of T cell memory in STING sufficient CD8 T cells. Thus, STING signaling regulates CD8 T cell memory fitness through both cell-intrinsic and extrinsic mechanisms. These studies provide insight into how IDO and STING therapies could improve long-term T cell protective immunity.

**Significance Statement:** STING signaling is an innate pathway that triggers host immunity against pathogens and cancer in response to cytosolic DNA. Additionally, STING signaling overactivation has been linked to autoimmunity. Yet, the interaction between antigenic and STING signaling and its impact in the development of protective immunity has remained unexplored. We found that strong levels of STING signaling impair CD8 T cell memory but only in response to high affinity TCR-pMHC interactions. Here, we provide evidence of how TCR signal strength controls STING signaling and IDO metabolism to regulate T cells’ survival as they mature to memory. These data have important implications for the design of STING and IDO combination immunotherapies

## Introduction

Innate and adaptive immunity are crucial to provide protection against infection. While innate immunity is the first arm of defense against the spread of pathogens, adaptive immunity relies on the activation of B and T lymphocytes with specialized functions to generate pathogen-specific memory cells able to provide long-term protection in the case of re-infection. Innate and adaptive immunity utilize different biochemical and signaling mechanisms to regulate immune cells’ responses. Growing evidence suggests that T cells express innate sensors(1), although whether these serve to fulfill a role in lymphocytes distinct from innate cells is still poorly understood.

Stimulator of IFN genes (STING) signaling is an innate sensor of pathogen and self-DNA that mediates inflammation(2). Upon infection, the synthase cGAS binds to double stranded DNA from viruses or bacteria and generates cyclic 2’3’GMP-AMP (cGAMP). ER-STING has high affinity for cGAMP and for bacterial cyclic dinucleotide(1). Upon binding, STING moves to the Golgi, and associates with TANK-binding kinase1 (TBK1) to facilitate IRF3 nuclear translocation and gene expression. Gain of function mutations in TMEM173 encoding STING have been associated to the autoinflammatory syndrome SAVI and are common in patients with systemic LUPUS(3–5). T cell cytopenia is a common feature of SAVI patients and recent reports using SAVI mouse models have concluded that the loss of T cells is IFN independent(5, 6). Meanwhile, induction of STING signaling is beneficial for T cell based anti-tumor immunity. This has triggered growing interest in developing STING agonists to improve cancer treatments(7). The role of STING signaling in T cell responses during infection, however, has remained largely unexplored.

Here, we have studied the impact of changing the levels of T cell extrinsic and intrinsic STING signaling in CD8 T cell responses to infection. Our data indicates that STING signaling plays a critical role in the development of T cell memory, and this role depends on TCR signal strength.

## Results

### Excessive STING signaling impairs CD8 T cell memory

*Listeria monocytogenes* (LM) is a model organism for the study of T cell responses and CD8 T cell differentiation in response to infection(8–12). The primary cytosolic sensor of *L. monocytogenes* is STING. Cyclic diadenosine monophosphate (c-di-AMP) secreted through bacterial multi-drug pumps triggers STING signaling in host cells and induces interferon responses(13, 14). Previous studies have shown that access to high levels of c-di-AMP can lead to a defect in cell mediated immunity(15). However, from these studies it was still unclear whether STING signaling affected T cell memory development or T cell memory responses or both.

To assess this, we performed adoptive transfer experiments using OT-1 CD8 T cells that specifically recognize the antigen OVA in the context of Kb. Then, we monitored OT-1 STING sufficient CD8 T cell differentiation during the immune response to a *L. monocytogenes* strain expressing the antigen SEINFEKL from ovalbumin/OVA (LM-OVA) or a variant of this strain that was engineered to hyper secrete the STING agonist c-di-AMP (referred here after as LM-OVA-STING^HI^ or STING^HI^ conditions)(15). We observed that the kinetics of the OT-1 T cell immune response were indistinguishable between STING sufficient and deficient hosts in wild type (WT) STING LM-OVA conditions (**Fig. 1B**). At the peak of the response, we also did not find any difference in the frequencies of OT-1 effector CD8 T cells in blood, suggesting CD8 T cell expansion was not impacted by increased or defective STING signaling. However, after the peak of the response the rate of contraction was increased in STING^HI^ conditions and almost no OT-1 CD8 T cells were detected 30 days post infection (**Fig. 1A-B**). Indeed, there was a dramatic reduction in the number of OT-1 memory T cells in both lymph nodes and spleen in STING^HI^ conditions, showing that excessive STING signaling leads to a defect in the establishment of T cell memory (**Fig. 1C**). Since enhanced STING signaling did not lead to an overt loss of CD4 or CD8 T cells, we concluded that the effect of STING signaling is restricted to antigen specific CD8 T cells (**Fig. 1D-E**).

**Figure 1.**
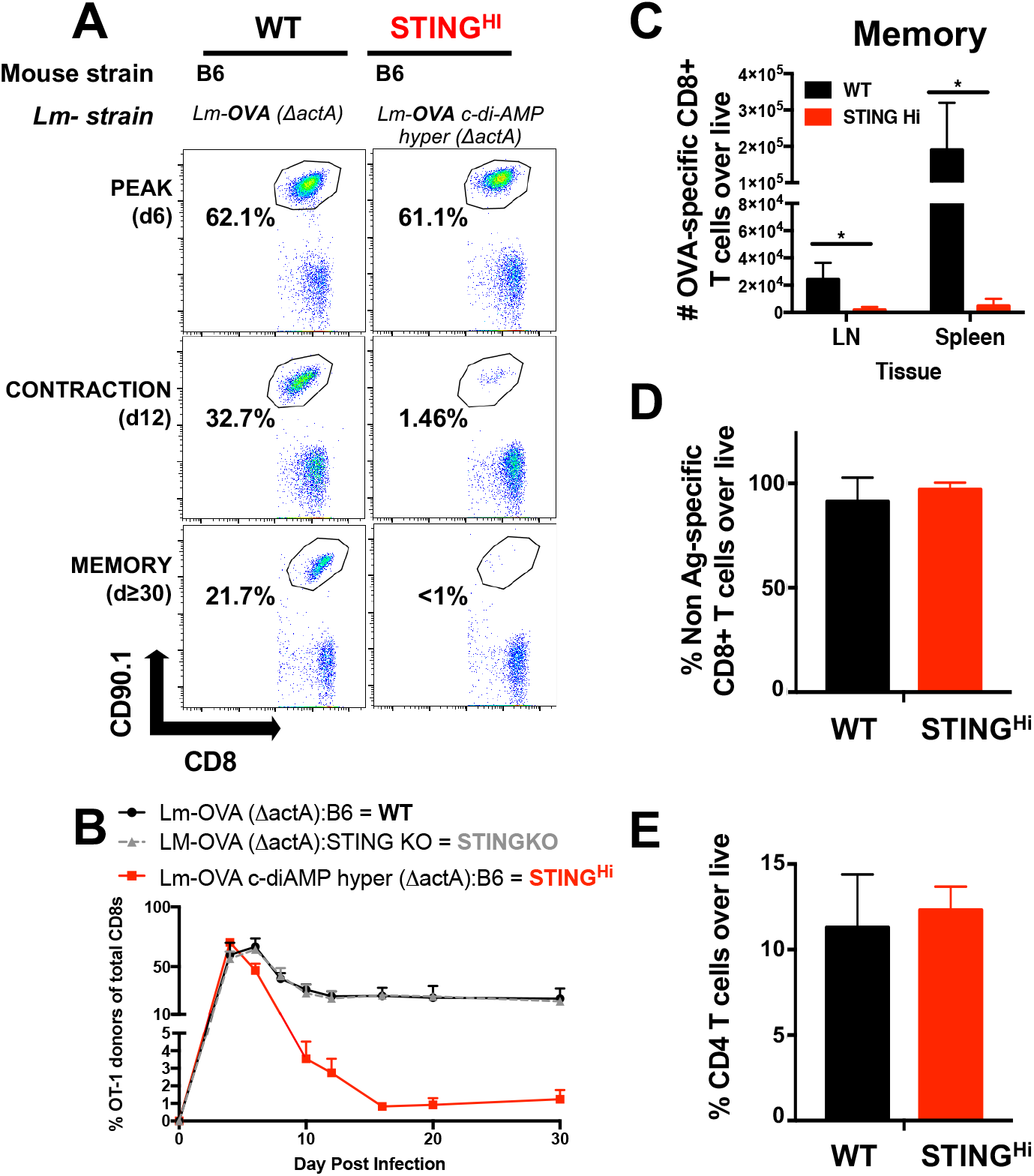
Excessive STING signaling impairs CD8 T cell memory. **(A-E).** Naïve OT-1CD90.1 T cells were transferred into congenic B6 hosts or Goldenticket (STINGKO) mice and challenged with LM-OVA ΔactA (WT) or LM-OVA ΔactA c-di-AMP hyper (STING^HI^). **(A).** Frequency of donor OT-1 cells at day 6, 12, and ≥30 p.i. Dot plots are representative of 3 independent experiments, with n≥3 mice per condition. **(B)**. Kinetics of the immune response of OT-1 donors upon infection with LM-OVA ΔactA (WT) or infection LM-OVA ΔactA c-di-AMP hyper secretion (STING^HI^) from blood. **(C).** Total number of OT-1 memory cells at day ≥30 p.i. in lymph nodes (LN) and Spleen. **(D-E)**. Frequency of endogenous CD8 T cells and CD4 T cells at day 12 p.i. from blood. Data were analyzed using multi t test and two-way ANOVA. Data are representative of 3 independent experiments, with n≥3 mice per condition. (*p<0.05, **p<0.005, ***p< 0.0005). **n.s**. non-significant.

### STING signaling does not affect T cell memory programming

We reasoned that enhanced STING signaling during priming could lead to an unbalanced generation of memory precursors (MPECS) and/or short-lived effectors (SLECs)(16). However, we did not observe any difference regardless of the level of STING signaling (**Fig. 2B**).

**Figure 2.**
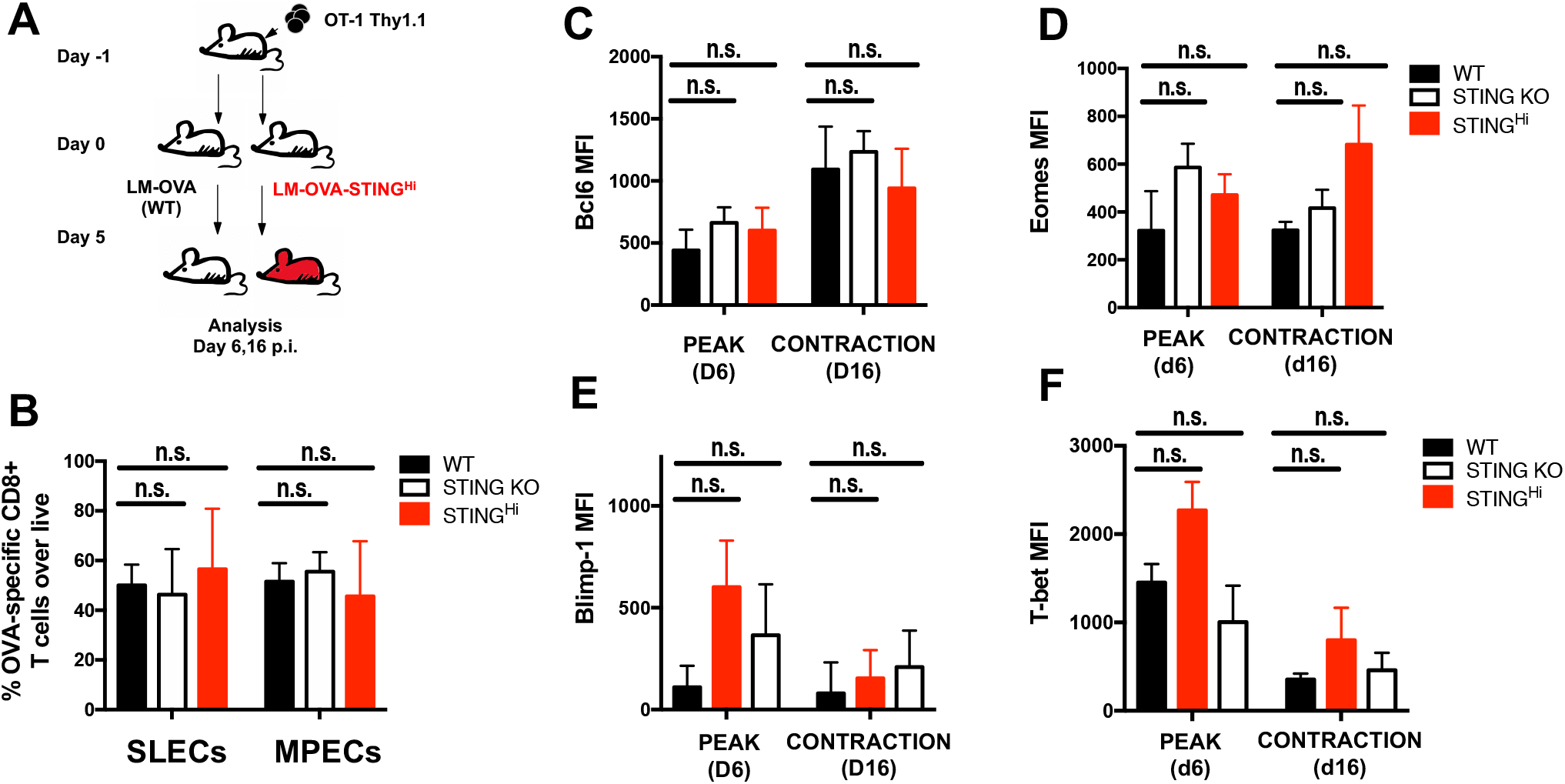
Enhanced STING signaling does not alter memory programming. **(A)** OT-1 naïve T cells were transferred into congenic hosts and challenged as described in Fig.1. **(B)**Frequency of OT-1 SLECs (CD127^lo^KLRG1^hi^) and MPECS (CD127^hi^KLRG^1lo^) at day 12 p.i. **(C-F)** Expression of Eomes, Bcl-6, T-bet, and Blimp-1 determined by flow cytometry from blood on OT-1 donor T cells. Data were analyzed using two-way ANOVA. Data obtained from at least 2 independent experiments, n≥3 mice. **n.s.** non-significant.

Next, we examined the expression of critical transcription factors for CD8 T cell memory programming, T-bet, Eomes, Blimp-1and Bcl-6 (17). Consistent with previous reports(17), as T cells progressed into their differentiation from the peak to the contraction phase of the response, their levels of Eomes and Bcl-6 memory associated transcription factors remained high, while T-bet and Blimp-1 effector associated transcription factors decreased (**Fig. 2C-F, Fig. S1**). This pattern of expression was the same in all conditions, indicating that enhanced STING signaling does not impair T cell memory programming.

### Increased STING signaling impairs the survival of CD8 T cell responders as they mature to memory

After the peak of a T cell immune response, most of the antigen specific CD8 responders die by apoptosis, with only 1-5% of them persisting to enter the memory pool(18). Since Bim, Bcl-xL and Bcl-2 are key for T cell survival during the contraction phase of the response(19–24), we asked whether STING signaling could affect their expression in CD8 T cells as they mature to memory. We found no differences in the expression of anti-apoptotic proteins Bcl-2 or Bcl-xL, neither at the peak nor at the contraction phase of the response. By contrast, Bim levels were much higher in CD8 T cells differentiating under enhanced STING signaling (**Fig. 3A-C**). This was observed both in SLECs and MPECs and was consistent with an increased in the frequency of Cleaved-caspase 3 positive antigen specific CD8 T cells under STING^HI^ conditions (**Fig. 3D-E**), indicating increased apoptosis.

**Figure 3.**
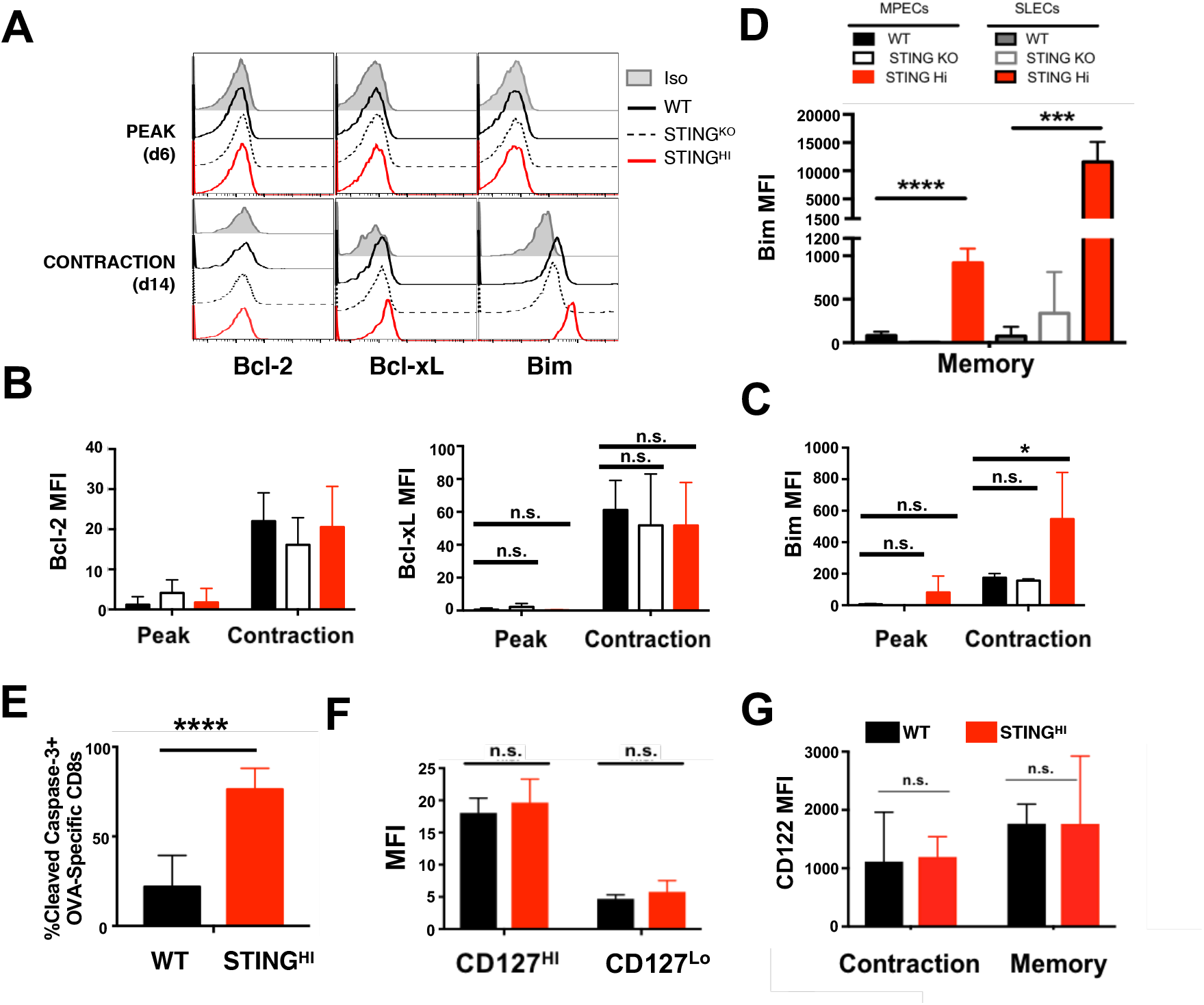
Enhanced STING signaling alters the apoptotic to survival balance in T cells maturing to memory. (**A-E)** OT-1 naïve T cells were transferred into congenic hosts and challenged as described in Fig.1. **(A-C)**. Expression of Bcl-2, Bcl-xL, and Bim determined by flow cytometry on OT-1 donor T cells at day 6 and day 14 p.i. Representative Histograms at day 6 and day 14 p.i. **(D)** Expression levels of Bim in SLECs and MPECs at d≥30 p.i. **(E)** Frequency of OT-1 donors positive for cleaved caspase-3. **(F-G).** Expression of CD127 (IL-7Rα) and CD122 (IL-2Rβ; IL-15R) surface levels at day 11(**F**) and day 11 and 30 p.i. (**G**). Data were analyzed using two-way ANOVA. Representative of 2 independent experiments, n≥3 mice per condition. (*p<0.05, **p<0.005, ***p< 0.0005), ****p<0.0001. **n.s.** non-significant.

We also did not find any change in the expression of receptors for homeostatic cytokines IL-7 and IL-5 associated with T cell memory survival(25), regardless of the levels of STING signaling (**Fig. 3 F-G**). Collectively, these data shows that an excess of STING signaling can cause increased apoptosis in antigen specific CD8 T cells, thereby, diminishing the generation of memory CD8 T cells.

### IDO metabolism regulates Bim levels and CD8 T cell memory under high levels of STING signaling

In the context of cancer and autoimmunity, STING signaling can induce IDO activity (26, 27) on innate cells and lead to a decrease in tryptophan levels that ultimately provoke T cell apoptosis(28). To test whether IDO metabolism could be involved in the mechanism by which enhanced STING signaling limits T cell memory, we adoptively transferred naïve OT-1 CD8 T cells into congenic hosts followed by infection with LM-OVA or LM-OVA-STING^HI^ and treated the mice with an IDO inhibitor, at the time where T cells are transitioning to memory (**Fig. 4A**). Treatment with the IDO inhibitor led to a recovery in T cell memory numbers that were reduced upon LM-OVA-STING^HI^ infection (**Fig. 4B-C**). Furthermore, we also found that the high Bim levels exhibited by antigen specific T cell responders under conditions of strong STING signaling returned to basal levels upon IDO inhibition. Bcl-2 levels remain similar to control and Bcl-xL levels only increased at memory (**Fig. 4D**). Importantly, IDO inhibition did not alter the formation of MPECs or SLECs (**Fig. 4E**), suggesting that the recovery of T cell memory is a result of impairing the Bim-dependent apoptosis driven by high levels of STING signaling.

**Figure 4.**
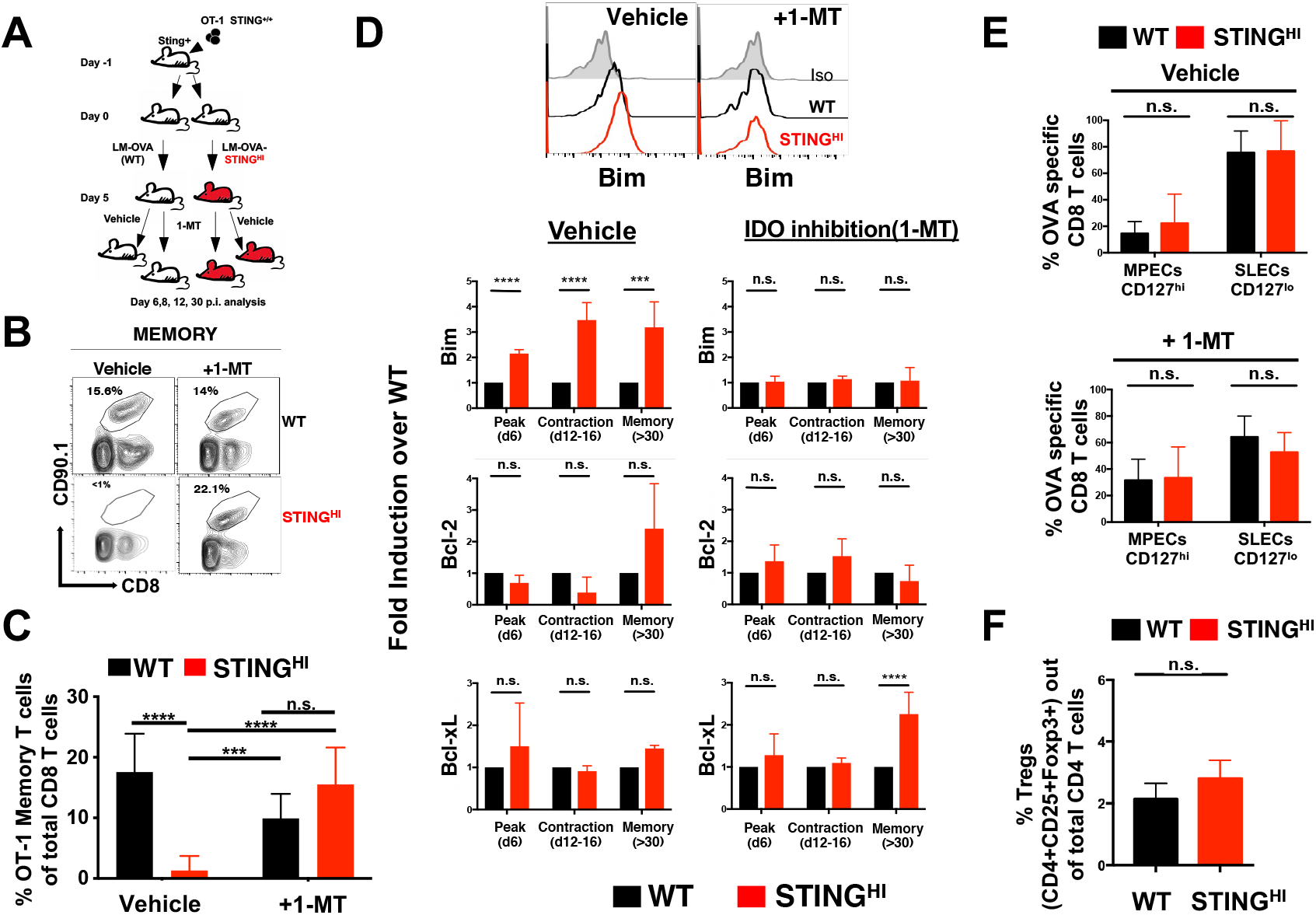
Inhibition of IDO restores CD8 T cell memory and Bim levels in T cells generated under enhanced STING ligands. **(A)** OT-1 naïve T cells were transferred into congenic hosts and challenged as described. Recipient mice were treated at day 5-30 p.i. with IDO inhibitor (+1-MT) or vehicle control. **(B)** Representative dot plots of donor T cells at day 30 p.i.. **(C)** Frequency of OT-1 cells with and without 1-MT treatment at memory (day ≥30 p.i.). **(D).** Bim, Bcl-2 and Bcl-xL expression over corresponding Vehicle and individual WT values through the course of the immune response in blood. **(E)** Frequency of OT-1 SLEC (CD127^lo^KLRG1^hi^) and MPEC (CD127^hi^KLRG^1l0^) donors at day 12 p.i. from blood. **(F)**. Frequency of regulatory T cells at day 8 p.i.. Data were analyzed using multi t test and two-way ANOVA. Representative of 2 independent experiments with n≥3 mice per condition. (*p<0.05, **p<0.005, ***p< 0.0005).**n.s.** non-significant.

Increased generation of regulatory CD4 T cells (Tregs) has been associated with IDO and tryptophan catabolism via the metabolite kynurenine(29-31). Yet we did not find any increase in Tregs under STING^HI^ conditions (**Fig. 4F**). Altogether, our results support a role for the IDO/STING axis in the regulation of CD8 T cell memory.

### T cell intrinsic STING signaling drives apoptosis of CD8 T cells differentiating to memory

Since both T and B cells can signal through STING(32–34), we also tested the contribution of T cell intrinsic STING signaling in the generation of CD8 T cell memory. For this, we adoptively transferred OT-1 STING sufficient (STING^+/+^) or deficient (STING^-/-^) CD8 naive T cells into STING sufficient congenic hosts, followed by infection with WT LM-OVA or LM-OVA-STING^HI^ (**Fig. 5A**). First, we evaluated the ability of antigen specific CD8 T cells to induce STING signaling *in vivo* during the context of infection. As expected, at day 6 p.i., we observed that OT-1 STING sufficient CD8 T cells exhibited higher levels of phosphorylated IRF3 upon LM-OVA-STING^HI^ than upon LM-OVA infection. By contrast, in STING deficient CD8 T cells, the induction of phosphorylated IRF3 was negligible (**Fig. 5B**). These data show that CD8 T cell responders can signal through STING *in vivo* in the context of infection.

**Figure 5.**
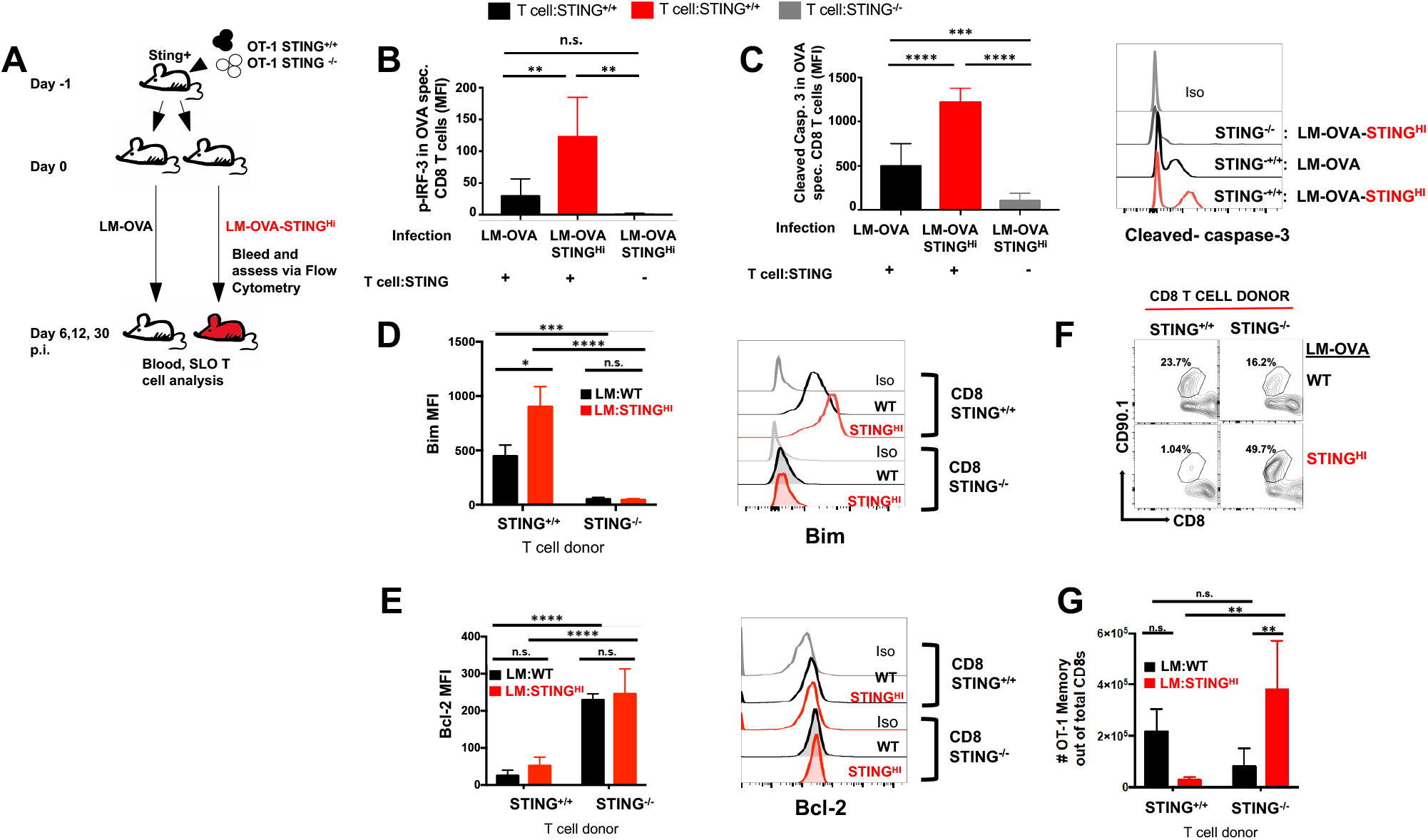
T cell intrinsic STING signaling drive IFN responses and T cell apoptosis during infection. **(A)** OT-1(STING^+^) or OT-1xSTING^-/-^ naïve T cells were transferred into STING sufficient congenic hosts and challenged with LM-OVA (WT) or LM-OVA-STING^HI^**. (B)** Expression of phosphorylated IRF-3 at day 6 p.i. in OT-1(STING^+^) or OT-1xSTING^-/-^ cells from blood. **(C)** Expression of Cleaved Caspase-3 at day 12 p.i. from blood. Representative histogram and graphs. **(D)** Representative histograms and graphs of the Expression of Bim **(D)** and Bcl2**(E**) at day 12 p.i.in blood. **(F).** Total numbers of OT-1 memory cells at day 30 p.i. isolated from LN. Data were analyzed using two-way ANOVA. Representative of 2 independent experiments. (*p<0.05, **p<0.005, ***p< 0.0005). **n.s.** non-significant.

Next, we measured the contribution of T cell-intrinsic STING signaling to T cell death during contraction. For this, we compared the levels of cleaved-Caspase-3(35), in STING sufficient or deficient T cells upon infection with the different LM-OVA strains. As expected, the levels of apoptosis (indicated by the levels of cleaved-caspase-3) increased as the levels of STING signaling augmented for WT T cells. This was not the case, however, for STING deficient T cells (**Fig. 5C**). Importantly, the reduction of apoptosis in STING^-/-^ T cells correlated with a reduction in Bim levels and an increase in Bcl-2 levels (**Fig. 5D-E**). Thus, T cell intrinsic STING regulates T cell survival during contraction.

Finally, we assessed the generation of T cell memory in T cells lacking STING. Upon LM-OVA-STING^HI^ infection, memory WT T cell numbers were extremely low, as expected (**Fig. 1, 5F**). In striking contrast, deletion of STING in CD8 T cell responders led to a recovery in T cell memory (**Fig. 5F-G)**. Collectively, these data are consistent with the idea that T cell intrinsic STING signaling regulates the rate of apoptosis of CD8 T responders during contraction and thereby, the establishment of CD8 T cell memory.

### TCR signal strength regulates the impact of STING signaling in CD8 T cell memory

CD8 T cell clones whose TCRs bind to antigen (peptide-MHC) with both high affinity and low affinity participate in the immune response and populate the T cell memory pool(36–38). Yet, while the memory pool holds an abundance of high-affinity T cell clones (due to their extraordinary proliferative rate), low-affinity T cell clones are preferentially programmed to become memory T cells(37). We evaluated whether the impact of STING signaling in the generation of CD8 T cell memory was dependent on the strength of antigenic signal. For this, we took advantage of a library of variants of the cognate antigen OVA previously characterized for their TCR affinity, potency, and ability to generate CD8 T cell memory in the OT-1 TCR transgenic model(36, 37, 39). We generated WT and supersecreting c-di-AMP *L. monocytogenes* strains that express low-affinity variants of OVA. We tested our hypothesis with LM-Q4H7 as it supports the generation of a discrete but detectable population of memory T cells (37, 39).

We repeated similar experiments to the ones in **Fig 1** using LM-Q4H7 and LM-Q4H7-STING^HI^ strains. As previously described by us and others(37, 39), the frequency of OT-1 T cells that were recruited to the peak of the response to LM-Q4H7 was smaller than upon LM-OVA infection due to low proliferation(36, 37). Yet we did not observe any difference in the frequency of antigen specific CD8 T cells generated in response to WT and STING^HI^ conditions, neither at the peak, contraction, or memory (**Fig. 6 A-C**). Indeed, the generation of memory T cells in response to LM-Q4H7-STING^HI^ was indistinguishable from the one in response to LM-Q4H7 (**Fig. 6C**). This contrasted with the dramatic loss of memory CD8 T cells that we observed in response to high STING signals and the high affinity antigen OVA (**Fig. 1**). Therefore, only in response to strong TCR signals, not weak, was T cell memory impaired by high STING signaling. Similar to what we observed in Fig. 1, survival of non-antigen specific CD8 or CD4 T cells was also not compromised (**Fig. S2**)

**Figure 6.**
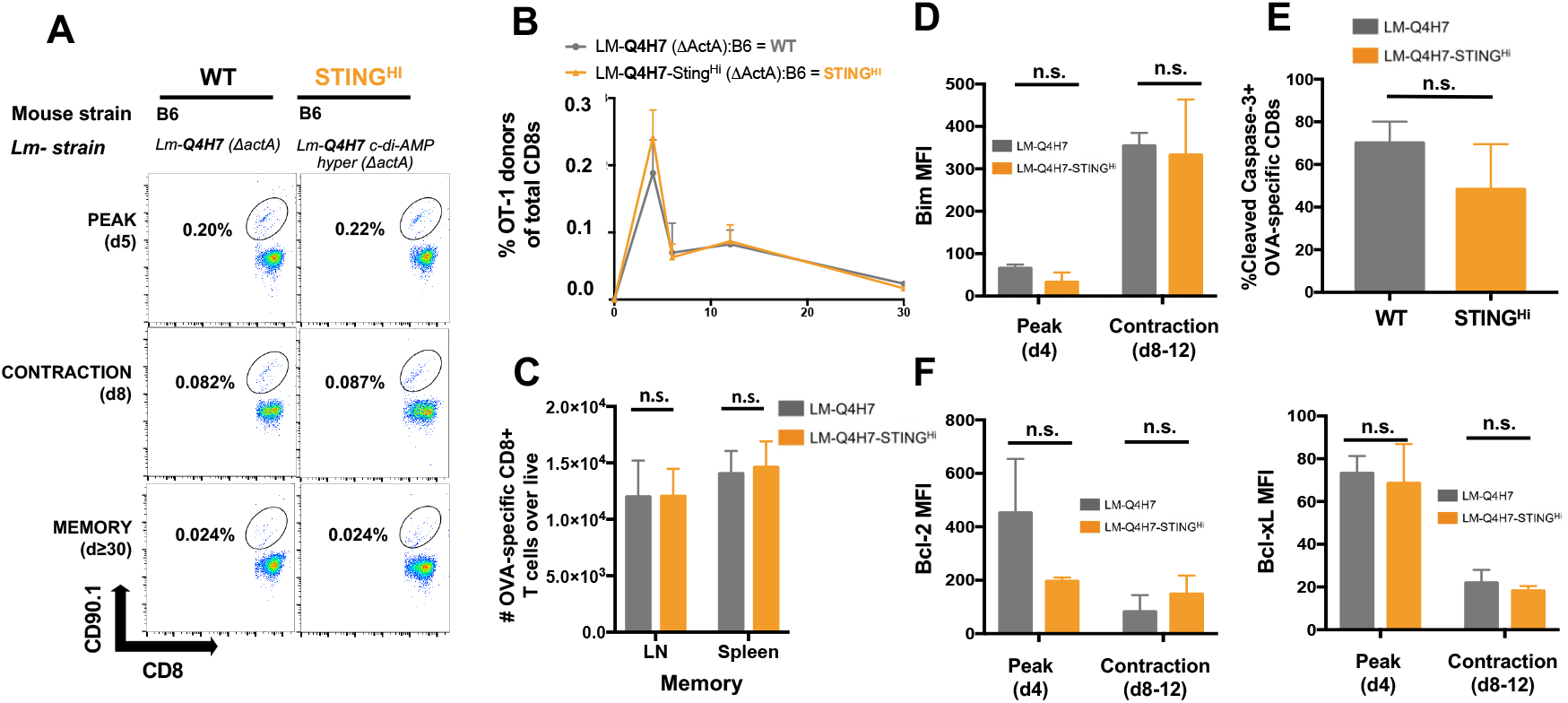
High levels of STING signaling impair CD8 T cell memory in response to strong but not weak TCR antigens. (**A-E)**. Naïve OT-1 T cells were transferred into congenic B6 hosts and challenged with LM-Q4H7 (WT) or LM-Q4H7-STING^Hi^. **(A)** Frequency of OT-1 cells at day 5, 8, and ≥30 p.i.. Dot plots are representative of 3 independent experiments, with n≥ 4 mice per condition. **(B)** Kinetics of the immune response of OT-1 T cells upon LM-Q4H7 ΔactA (WT) in B6 congenic host, gray line; or upon infection with parental c-di-AMP hypersecreting LM-Q4H7 ΔactA strain in B6 congenic host (STING^HI^), gold line. **(C)** Total number of memory donor OT-1 cells at day 32 p.i.. Expression of Bim (**D**), Bcl-2**(F),** and Bcl-xL (**G**). **(E)** Frequency of cleaved caspase-3 positive OT-1 T cells at day 12 p.i. Data were analyzed using multi t test and two-way ANOVA. Data are representative of 3 independent experiments, with n≥ 4 mice per condition. (*p<0.05, **p<0.005, ***p< 0.0005). **n.s.** non-significant.

CD8 T cells responding to LM-Q4H7-STING^HI^ exhibited levels of Bim and apoptosis (cleaved caspase-3) as well as pro-survival factors Bcl-2 and BclxL comparable to controls (**Fig. 6 D-F**). Furthermore, we did not find any difference in the expression of IL-7 and IL-15 receptors, which are important for T cell memory survival(25) (**Fig. S3**). Importantly, increased STING signaling did not lead to further skewing towards T cell memory, as there were no differences in SLECs, MPECs or memory associated transcription factors between T cells responders to LM-Q4H7 or LM-Q4H7-STING^HI^. (**Fig. S4**).

Finally, we tested whether enhancing STING signaling during priming would confer any advantage to the response of low-affinity memory T cells to challenge. We immunized mice carrying the same number of OT-1 donors with LM-OVA, LM-OVA-STING^HI^, LM-Q4H7 and LM-Q4H7-STING^HI^. At day 30 p.i. half of the cohorts were challenged with WT-LM-OVA and the other half sacrificed to determine the generation of primary OT-1 T cell memory. Then, we compared the generation of primary and secondary OT-1 T cell memory among all conditions. As expected, the frequency of OT-1 memory T cells primed with LM-OVA increased upon challenge. Secondary memory was considerably increased (almost to the LM-OVA control levels) for the LM-Q4H7 and LM-Q4H7-STING^HI^ conditions. The increase in secondary memory after LM-OVA-STING^HI^ immunization and LM-OVA challenge, however, was minimal (**Fig. S5A**). These data suggest that CD8 memory T cells generated in response to both strong TCR and STING signals are impaired in their recall response, while this is not the case for T cells responding to weak TCR signals or weak antigens.

To further test the CD8 T cell memory response, we compared the effector function of memory CD8 T cells generated upon the different immunizations in the context of cancer. For this, we generated OT-1 memory CD8 T cells upon immunization with LM-OVA, LM-Q4H7 and LM-Q4H7-STING^HI^ (The number of OT-1 memory generated upon LM-OVA-STING^HI^ was too low to perform the experiment). At day 30 p.i, we harvested OT-1 memory CD8 T cells and transferred them in equal numbers into hosts that had been previously implanted with OVA expressing tumor cells. Then, we monitored tumor growth to assess the ability of memory OT-1 T cells generated in the different antigenic and STING conditions to control tumors. High affinity (LM-OVA primed) memory T cells controled tumor growth better than non-treated controls, as expected (**Fig. S5 B-C**). Most importantly, there was a significant delay in tumor growth in mice treated with low affinity memory CD8 T cells generated under strong STING signaling (**Fig. S5 B-C**). Low affinity memory T cells generated under WT STING levels controlled the tumor better early than high affinity memory T cells. However, at later time points tumor growth began to rebound. By contrast, low affinity memory T cells generated under strong STING signals, controlled tumor burden for longer. These data suggest that while enhanced STING signaling does not impact the generation of low affinity memory CD8 T cells, it improves T cell memory responses. Thus, the impact of STING signaling in the generation and response of memory CD8 T cells depends on TCR signaling strength during priming.

### TCR signal strength controls STING levels and ER-stress dependent proteins associated with apoptosis

Given the connection that we found between TCR signal strength and STING signaling, we explored whether T cells activated by high affinity (OT-1:OVA) and low affinity (OT-1:Q4H7) TCR ligands induce similar STING signals. We stimulated high and low affinity CD8 effector T cells with STING agonists in the presence or absence of further TCR stimulation. We found that stimulation with STING agonists induce a lower level of pIRF3 in low than high affinity CD8 T cells. This correlated with a lower level of phosphorylated STING (higher MW)(40) (**Fig.7A-B)**, as well as lower levels of Bim and cleaved-caspase-3 (**Fig 7A-D**). Therefore, T cells stimulated by weak TCR signals or low affinity antigens are impaired in STING signaling, Bim expression and apoptosis.

**Figure 7.**
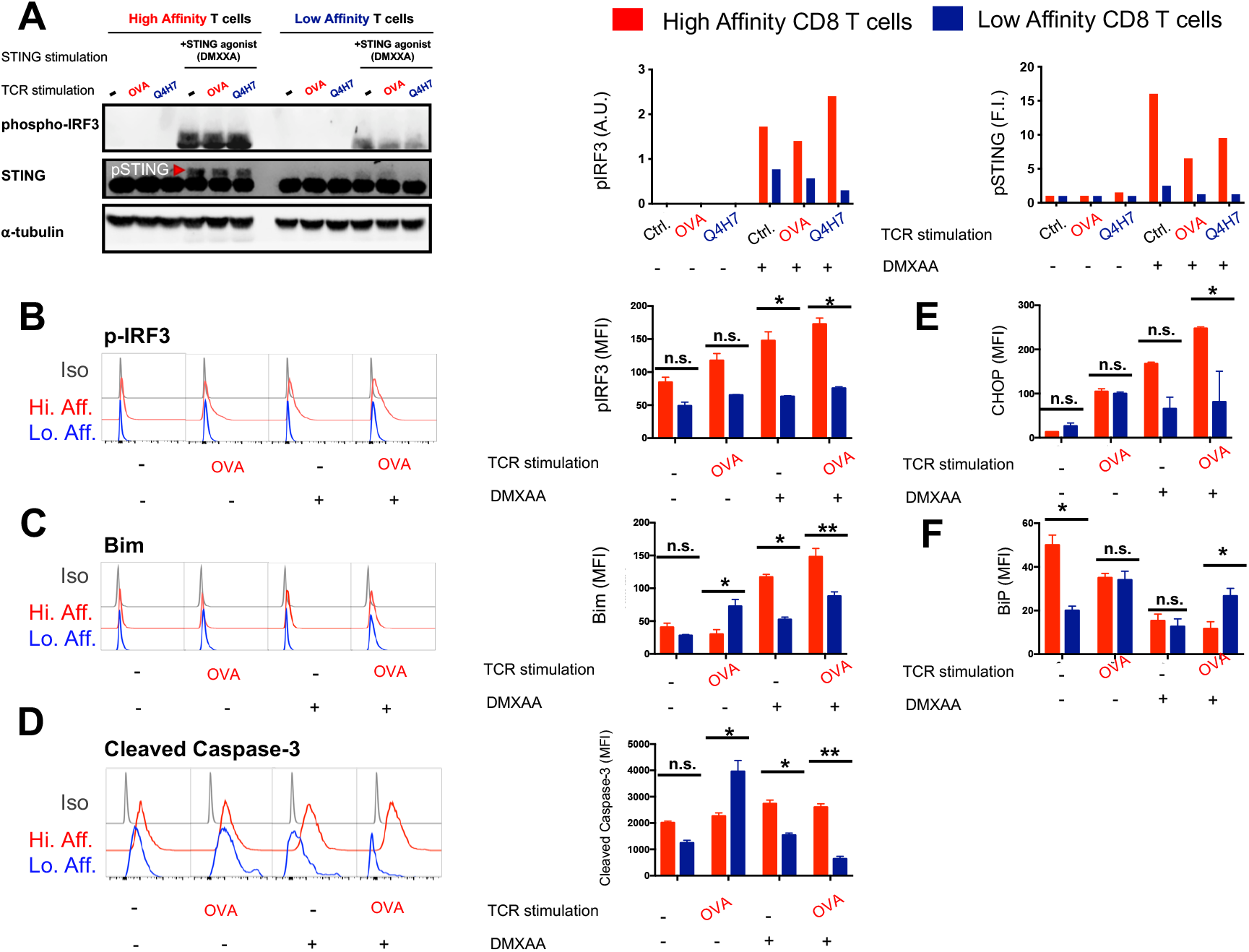
TCR signal strength controls the threshold of STING signaling that causes T cell apoptosis via the UPR pathway. (**A-C**). Naïve OT-1 splenocytes were stimulated with OVA or Q4H7 peptides as in (64). At day 3 post stimulation, OT-1 cells were restimulated for 6 hours with either Kb-OVAtet or Kb-Q4H7tetramers as in (65) in the presence or absence of STING agonists. **(A)** Whole cell lysates were resolved in SDS-PAGE and the expression of proteins indicated (phosphorylated IRF3 and STING) was determined by immunoblot. **(B)** Representative histograms of p-IRF3, Bim and cleaved caspase-3 in high and low affinity T cells (left) and their bar graphs (**C)**. **(E-F)** Bar graphs showed expression of ER proteins CHOP and Bip. Cells were stained for flow cytometry and indicated protein MFIs were calculated by subtracting Isotype controls. Data are representative of 2 independent experiments, with n≥ 3 per condition, and were analyzed using two-way ANOVA. (*p<0.05, **p<0.005, ***p< 0.0005). **n.s.** non-significant.

Emerging evidence suggests that ER stress associated with STING activation can elicit apoptosis(41, 42). Under cell stress, the Unfolded Protein Response (UPR) is triggered and chaperone protein Bip dissociates from stress sensors anchored in the ER membrane, leading to the expression of the transcription factor CHOP. CHOP in turn, increases the expression of Bim resulting in cell apoptosis(43-45). Upon stimulation with STING agonists, we found that low affinity T cells expressed significantly lower levels of CHOP than high affinity T cells (**Fig. 7E and Fig. S6**). Thus, weak TCR signals pose T cells less sensitive to STING activation, which results in lower expression of the ER-stress protein CHOP, Bim and apoptosis. This further suggests that the strength of antigenic signals control the threshold of STING signaling that causes T cell apoptosis via the UPR pathway..

## Discussion

We have found that STING signaling can severely impair the generation of memory CD8 T cells during infection, depending both on the level of STING and antigenic signaling. T cells express STING and TCR signals control the ability of T cells to be resistant to apoptosis-mediated STING signals.

As an innate sensor, STING’ s presence in lymphocytes could be interpreted as an ancestral mechanism that enables most of the cells in the body to respond to infection by default. STING pathway is, indeed, highly conserved through evolution(46) and T cells, themselves, can trigger the IFN response upon STING ligand stimulation(32). However, the preferred response of T and B lymphocytes to STING agonists is to die(32, 47). This raises the question as whether STING could serve other purposes in T cells beyond IFN responses. The massive T cell death observed after the peak of an immune response aids in regaining organismal homeostasis. Thus, it is possible that during infection T cell intrinsic STING signals function to maintain cell homeostasis. STING-dependent T cell death would not only regulate T cell numbers but would also impair their potential to induce excessive IFN responses.

Although our data strongly suggest that the level of TCR signaling at priming predisposes effector CD8 T cells to trigger weaker STING signaling in response to STING agonists, how this occurs is unclear. A connection between the BCR, MHC receptors and STING has been previously shown. BCR and STING signaling appeared to regulate both BCR signaling and BCR downregulation(34, 48), however this seems to be a synergistic interaction and not a multistep process as it is deduced from our T cell studies(49, 50). On the other hand, MHC class II proteins can associate with MYPs or STING and mediate transduction of apoptotic signals(33). Could the TCR operate in a similar fashion? A recent report showed a costimulatory effect between TCR and STING signals that serves to sustain IFN responses in activated CD4 T cells (51) but a connection to T cell apoptosis was not investigated. The studies presented here show a link between TCR signal strength and STING-dependent T cell survival. It is tempting to speculate that membrane proximal events similar to the ones that occur in B cells and APCs might operate in T cells (similar to B and APCs) to allow a crosstalk of STING and TCR signals(33, 50).

We have found that T cell intrinsic and extrinsic STING mechanisms contribute to the establishment of T cell memory. T cell extrinsic STING mechanisms usually imply the activation of STING in dendritic cells and induction of costimulatory molecules that further promote T cell activation(52). Our results point at IDO as a critical mediator of T cell death during the contraction phase of the immune response, which limits CD8 T cell memory. In the context of cancer, cell death activates STING signaling resulting in expression of type I IFN. This leads to an induction in IDO expression on innate cells to promote Trp catabolism. Trp catabolism, in turn, supports tumor growth and immune tolerance(53). In our model of infection, inhibition of IDO reverted the effects of STING signaling on proapoptotic-Bim expression. Interestingly, limited access to Trp (due to IDO activity) can also induce the ER stress response(54). Indeed, upregulation of the ER-stress protein CHOP has been observed in T cells exposed to conditions of tryptophan deprivation via IDO-expressing DCs(54, 55). Since TCR signaling also regulates STING-dependent induction of CHOP and CHOP, mediates Bim associated apoptosis(56), we propose that high levels of STING signaling causes ER-stress in T cells that result in the activation of the Unfolded protein response (UPR). Thus, both T cell extrinsic and intrinsic-STING mechanisms may converge at the level of the UPR and regulate the expression of CHOP-mediated apoptosis in T cells, controlling their survival as they mature to become memory T cells.

There is a great effort in developing STING agonists and inhibitors for therapy against cancer and autoimmune or inflammatory diseases such as LUPUS and SAVI(7). Yet, our understanding of the ramifications derived from changing the levels of STING signaling (either increasing it or inhibiting it) on inflammation and long-term adaptive immunity is very limited. Our study supports the idea that STING agonists dosage may need to be carefully calibrated to maximize anti-tumor immunity depending on the nature of the tumor antigens. Thus, for tumors that present neoantigens, excessive STING signaling may be detrimental while beneficial for tumors where self-tumor antigens are dominant. While this hypothesis will need to be further tested, a recent report found that lower doses of STING agonist regimens elicited better CD8 T cell antitumor responses than the tumor ablative higher doses(57). In our system, IDO inhibition counterbalanced the harmful effects of excessive STING signaling on the establishment of long-term T cell memory. Considering this, the detrimental effect of Trp metabolism in tumors(53) and the fact that IDO inhibitors are under preclinical and clinical evaluation for the treatment of cancer, we concur with others that regimens of STING agonists and IDO inhibitors may be more effective to generate long-term protection against tumor recurrence(58).

## Materials and Methods

### Mice and reagents

C57BL/6J (B6), B6.SJL-Ptprc^a^ Pepc^b^/BoyJ (CD45.1^+^ congenic C57BL/6), B6.PL (CD90.1^+^) OT-1, Goldenticket (Tmem173^gt^, STINGKO), and OT-1xGoldenticket (OT-1xTmem173^gt^, OT-1xSTINGKO)(59) mice were bred and maintained in accordance with University of Missouri Office of Animal Resources Animal Care and Use committee. Infection and maintenance of infected mice occurred in an ABSL2 facility at the University of Missouri. OVA (SIINFEKL), Q4H7 (SIIQFEHL), and VSV NP52-59 (RGYVYQGL) peptides were from New England Peptides. Biotinylated H-2K^b^ monomers were generated in our laboratory, and tetramerized using fluorescently tagged streptavidin from Biolegend. 5,6-Dimethylxanthenone-4-acetic Acid (DMXAA) was from Millipore Sigma. 1-Methyl Tryptophan (1MT) was from Sigma Aldrich.

### Antibodies and Flow Cytometry

Single-cell suspensions of tissues were labeled with antibodies for 30 min on ice. The BD Cytofix/Cytoperm (Cat#554714) was used to prepare the cells for intracellular staining per manufacturer’s instructions. Flow cytometry was performed on an LSRII Fortessa flow cytometer (BD) and analyzed with Flowjo FACS Analysis Software (Tree Star, Inc.) using the specific antibodies (SI appendix).

### Adoptive transfer

Naïve CD90.1 OT-1 CD8 T cells (1×10^4^) were adoptively transferred intravenously into C57BL/6 host mice. Memory OT-1 CD8 T cells (1×10^4^) were purified from the lymph nodes and spleen of previously immunized host mice carrying OT-1 memory T cells as CD90.1^+^CD8^+^ by magnetic sorting (see below).

### Magnetic bead purification

Before transfer into recipient mice, both naïve and memory OT-1 CD8 T cell donors were magnetically sorted via magnetic beads (Miltenyi Biotec) with specific antibodies. Naïve donors were sorted via negative selection (isolated as CD4^-^CD19^-^NK1.1^-^CD11c^-^) and memory OT-1 CD8 T cells were sorted via positive selection (isolated as CD8+CD90.1^+^) from host mice at day ≥30 post infection. A 100-90% enrichment after sorting, was verified in the naïve and memory OT-1 donor populations before adoptive transfer via flow cytometry.

### Bacterial Strains and Infections

Attenuated *Listeria* strains expressing either OVA (SIINFEKL) or OVA variant Q4H7 (SIIQFEHL) peptides with or without hyper cyclic-di-AMP production (LM-OVA ΔactA; LM-Q4H7 ΔactA; LM-OVA-STING^Hi^ ΔactA; LM-Q4H7-STING^Hi^ ΔactA) were constructed as follows. First, WT OVA (SIINFEKL) and Q4H7 (SIIQFEHL) OVA alleles^36^ were cloned under control of the *actA* promoter and in-frame with the ActA signal sequence in a derivate of pPL2 and integrated at the *tRNA^Arg^* locus(60) in the attenuated *ΔactA* background (Lm *ΔactA ΔinlB*)(*61*). Second, to generate the STING^Hi^ strains, *tetR*::Tn917 was transduced into each strain as described(15). *Listeria* strains were grown to an OD_600_ of 0.1 (as they enter log phase) in BHI broth. Pre- and post-injection *Listeria* CFUs were determined as in(62). 24hrs after OT-1 donor cells adoptive transfer, recipient mice received intravenous injections of 1×10^5^ CFU.

### Administration of 1-MT

IDO inhibitor 1-MT was provided in drinking water. 1-MT (Sigma-Aldrich D-1MT, catalog no. 452483) was prepared at 2mg/mL (pH 7-7.3). Before being delivered to the mice, the water was filter sterilized and supplemented with 1% sucrose to improve acceptance by the mice. Mice drank 4-5 mL/day, similar to consumption of water without drug. 1-MT was administered in the drinking water for 2-3 weeks, starting at day 5 post infection (freshly made 1-MT water was given every 5 days).

### In vitro Stimulation

Splenic naïve T cells were activated at day 0 at a concentration of 2×10^6^ cells/mL using OVA or Q4H7 peptides as in (37). Effector CD8 T cells were then restimulated at day 3 using Kb-OVA-or Q4H7-tetramers(39) and anti-CD28 (10μg/mL) with or without STING agonist DMXAA (10ug/mL) for 6hrs. T cells were then collected and subject to immunoblot. Tetramer concentration was adjusted as in (39)to achieve equal T cell occupancy for each ligand.

### Immunoblot analysis

In vitro stimulated T cells were harvested and lysed in buffer containing 20mM Tris/HCl pH 7.6, 150mM NaCl, 0.1mM EDTA pH 8.0, 0.1mM EGTA pH 8.0, 1mM β-glycerphosphate, 1% NP-40, 1mM Na_3_VO^4^, 1μg/ml Leupeptin, 1μg/ml Aprotinin, 1μg/ml Pepstatin, 1mM NaF, 1 mM PMSF. Samples were resolved on a 10% SDS-PAGE gel and transferred to a nitrocellulose membrane. Membranes were blocked with 5% nonfat dry milk (Bio-Rad) and probed using specific antibodies (SI appendix) according to the manufacture recommendations. Blots were imaged on a Li-Cor Odyssey XF and densitometry was analyzed and quantified using Image Studio program (Li-Cor).

### In vivo Tumor analysis

Congenic CD90.1^+^CD8^+^ OT-1 memory T cells were purified from adoptively transferred; infected mice after 30 days as described above. The sorted cells were then injected intravenously into B6 Rag-/-mice preciously inoculated with 5×10^5^ EG.7-OVA thymomas (6 days prior). Tumor area was measured every 2 days for 24 days as in (63).

## Acknowledgments

We thank Dr. Daniel Portnoy (UC Berkeley) for insightful discussion. We thank Samantha Hopps for assistance with screening and mouse work. MU Office of Research and School of Medicine animal vivarium staff for assistance with mice. Kathy Schreiber and Daniel Jackson for assistance in the Flow Cytometry Core Facility. Supported by National Institutes of Health grants R01 AI110420-01A1 and R56 AI110420-06A1, 1U01CA244314 as well as Internal funding from the School of Medicine (ET).

