## Supplemental methods and figures for "STING controls T cell memory fitness during infection through T cell intrinsic and IDO dependent mechanisms"

**Supplemental appendix**

| Antibodies for Flow Cytometry | | | |
|---|---|---|---|
| **Antigen** | **Clone** | **Fluorochromes** | **Vendor** |
| CD8 | 53-6.7 | BV785 | BD Biosciences |
| CD4 | L3T4 | APC | Biolegend |
| CD45.2 | 104 | BV605 | Biolegend |
| CD90.1 | OX-7 | PE-Cy7, BV605 | BD Biosciences |
| IL-2 | JES6-5H4 | PE | BD Biosciences |
| TNFα | MP6-XT22 | BV785 | Biolegend |
| IFN-g | XMG1.2 | BV421 | BD Biosciences |
| Granzyme B | GB12 | APC | Invitrogen |
| CD127 | A7R34 | FITC | Biolegend |
| CD122 | TM-b1 | PE | BD Biosciences |
| KLRG1 | 2F1 | APC | Biolegend |
| Bcl6 | BCL-DWN | PE | Invitrogen |
| Blimp1 | 5E7 | APC | BD Biosciences |
| Tbet | eBio4BIO | AF660 | Invitrogen |
| Eomes | Dan11mag | AF660 | Invitrogen |
| CD25 | PC61.5 | PE | Invitrogen |
| STING | D2P2F | Unconjugated | Cell Signaling |
| pIRF3 | D6O1M | FITC | Cell Signaling |
| Bim | C34C5 | Unconjugated | Cell Signaling |
| Bcl-2 | 3F11 | PE | BD Pharmingen |
| Bcl-xL | 7B2.5 | FITC | Southern Biotech |
| Cleaved Caspase-3 | D3E9 | APC | Cell Signaling |
| BiP | C50B12 | PE | Cell Signaling |
| CHOP | L63F7 | FITC | Cell Signaling |
| αRabbit Secondary | A11034 | AF488 | Life Technologies |
| αRabbit Secondary | A21429 | AF555 | Life Technologies |
| Vα2 | B20.1 | APC | Biolegend |
| Vβ5 | MR9-4 | PE | BD Pharmingan |
| CD62L | MEL-14 | PE-Cy7, BV785 | Biolegend |
| CD44 | IM7 | BV605 | Biolegend |

| Antibodies for Western Blotting | | | |
|---|---|---|---|
| **Antigen** | **Clone** | **Host Species** | **Vendor** |
| CD28 | 37.51 | Goat mAb | BD Biosciences |
| α-Tubulin | DM1A | Mouse mAb | Sigma |
| STING | D2P2F | Rabbit mAb | Cell Signaling |
| pIRF3 | D6O1M | Rabbit mAb | Cell Signaling |
| Goat anti-mouse | | Goat Polyclonal | Li-Cor |
| Goat anti-rabbit | | Goat Polyclonal | Li-Cor |

### SUPPLEMENTAL FIGURES

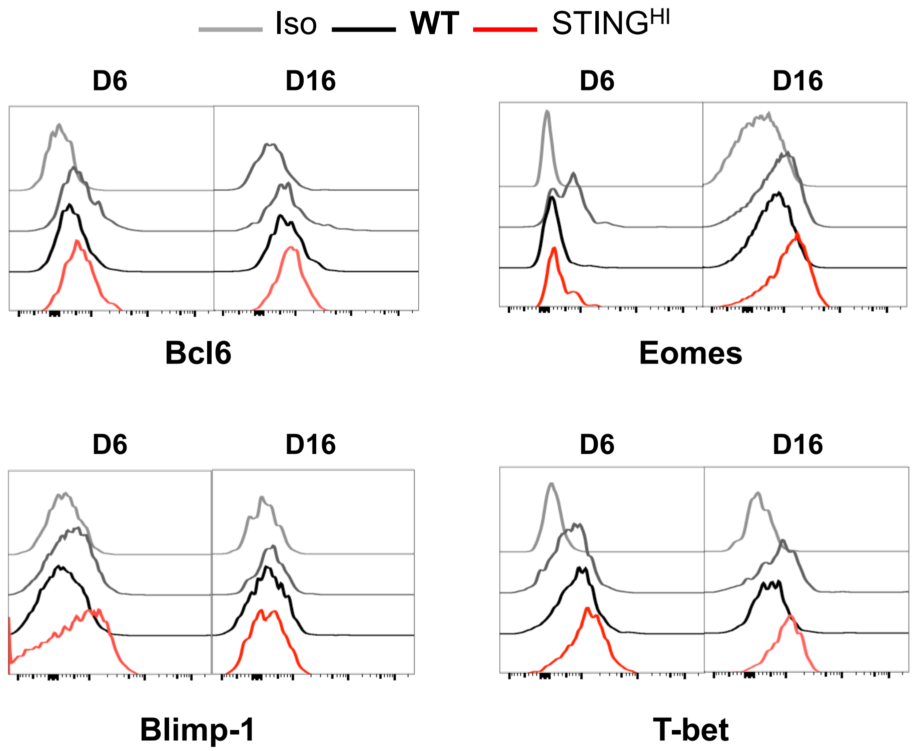


**Fig. S1.** Representative Histograms of the expression of transcription factors Bcl-6, Blimp-1, Eomes and T-bet at day 6 and 16 p.i. in donor OT-1 T cells responding to WT LM-OVA and WTLM-OVA-STINGhi. Linked to Figure 2.

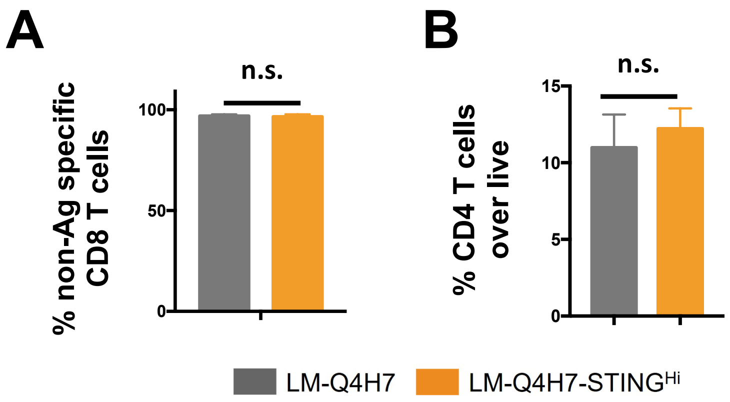


**Fig. S2. Hig levels of STING signaling do not impact the survival of non-antigen specific endogenous CD8 T cells nor CD4 T cells in response to weak TCR stimulation.** OT-1 naïve T cells were transferred into congenic hosts and challenged as described in **Fig. 6.** (**a-b)** Frequency of non-specific endogenous CD8 T cells and CD4 T cells at day 12 p.i. Data was analyzed using two-tailed t test and are representative of 3 independent experiments, with n$\geq$ 3 mice per condition. (*p<0.05, **p<0.005, ***p< 0.0005). **n.s**. non-significant. Linked to Fig. 6

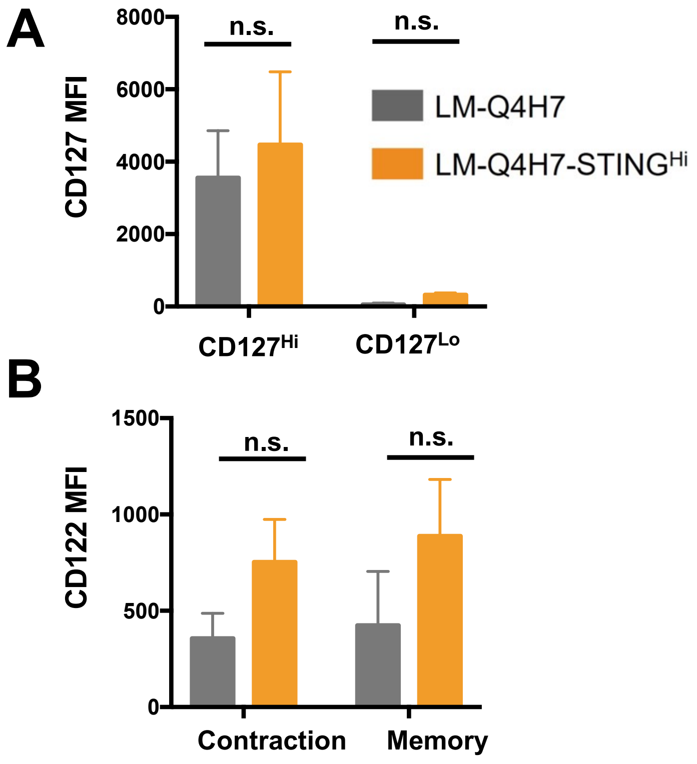


**Fig. S3. Enhanced STING signaling does not affect the expression of receptors for IL-7 and IL-15 in low affinity CD8 T cells.** OT-1 naïve T cells were transferred into congenic hosts and challenged as described in **Fig. 6. (a-b)** IL-7R (CD127) and IL-15R (CD122) surface levels at days 11 and 30 p.i. respectively. MFI, means fluorescence intensity. The data was analyzed using two way analysis of variance..(*p<0.05, **p<0.005, ***p< 0.0005). **n.s.** non-significant.

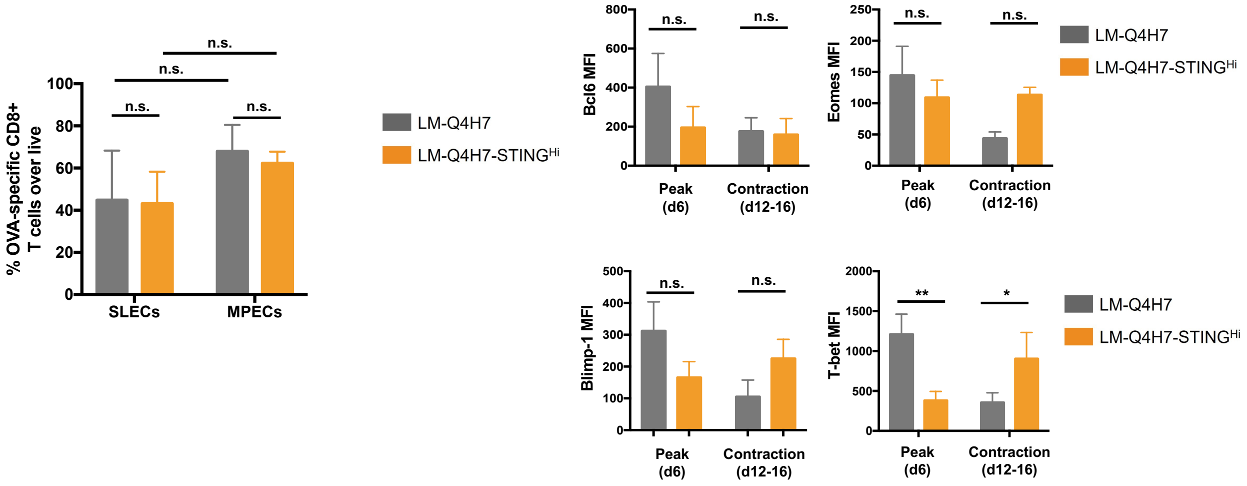


**Fig. S4. Enhanced STING signaling does not impact T cell memory programming in response to weak TCR stimulation.** OT-1 naïve T cells were transferred into congenic hosts and challenged as described in **Fig. 6. A.** Frequency of OT-1 SLECs (CD127$^{lo}$KLRG1$^{hi}$) and MPECS (CD127$^{hi}$KLRG$^{1lo}$) at day 12 p.i. **B.** Expression of Eomes, Bcl-6, T-bet, and Blimp-1 determined by flow cytometry from blood on OT-1 donor T cells at days indicated. MFI, means fluorescence intensity. The data was analyzed using two way analysis of variance. (*p<0.05, **p<0.005, ***p< 0.0005). n.s. non significant

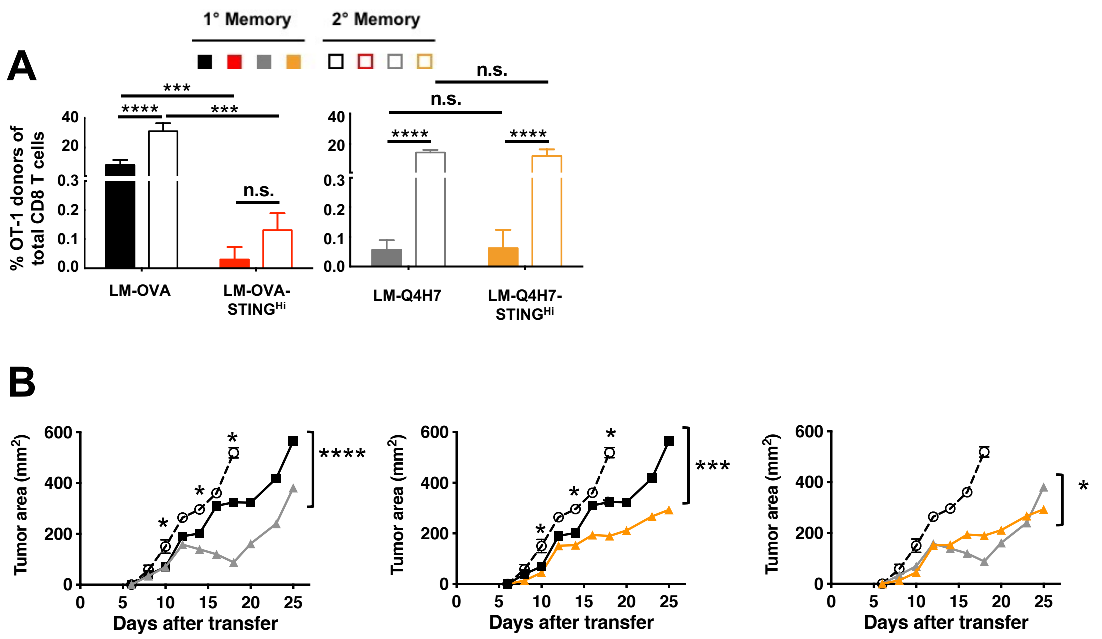


**Fig. S5. TCR signal strength regulates the impact of STING signaling in CD8 T cell memory responses. (a).** OT-1 naïve cells were transferred into B6 congenic hosts and challenged as described in **Fig. 1** in addition to LM-Q4H7 or LM-Q4H7-STING$^{hi}$. Graphs are representative of 3 independent experiments, with n≥3 mice per condition. OT-1 frequencies at memory in blood. Frequency of $1^0$ and $2^0$ memory OT-1 cells after re-challenge with LM-OVA (1x $10^5$ CFUs). **(B)** Memory T cells generated as in **(A)** were MACS purified at day 40 from spleen and lymph nodes and transferred in equal numbers into congenic hosts bearing EG7 thymomas. Graphs show tumor area over time after memory transfer. Representative of 2 independent experiments with n≥5 mice per condition. Data was analyzed using unpaired t tests fro a) Data was analyzed using a 2 way ANOVA for all curves with exception of multi t test for specific points between NT (clear) and LM-OVA (black). All differences between NT, Q4H7, and Q4H7-STING$^{hi}$ are significant. (*p<0.05, **p<0.005, ***p< 0.0005, ****p< 0.0001). **n.s.** non-significant.

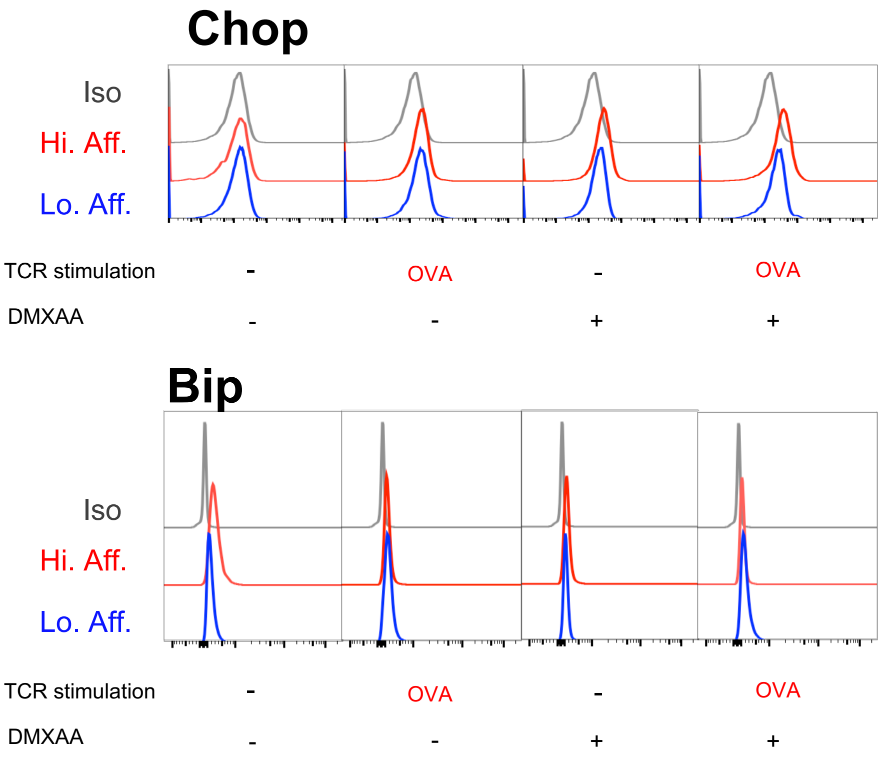


**Fig. S6**. Representative histograms for Chop and Bip expression in high affinity (red) and low affinity (blue) CD8 T cells upon stimulation with STING agonists. Grey histogram refers to isotype control. Data linked to Fig. 7
